# Insuperable problems of the genetic code initially emerging in an RNA World

**DOI:** 10.1101/140657

**Authors:** Peter R Wills, Charles W Carter

## Abstract

Differential equations for error-prone information transfer (template replication, transcription or translation) are developed in order to consider, within the theory of autocatalysis, the advent of coded protein synthesis. Variations of these equations furnish a basis for comparing the plausibility of contrasting scenarios for the emergence of tRNA aminoacylation, ultimately by enzymes, and the relationship of this process with the origin of the universal system of molecular biological information processing embodied in the Central Dogma. The hypothetical RNA World does not furnish an adequate basis for explaining how this system came into being, but principles of self-organisation that transcend Darwinian natural selection furnish an unexpectedly robust basis for a rapid, concerted transition to genetic coding from a peptide•RNA world.

## 1 Introduction

The RNA World (Gilbert 1986) is a widely-embraced hypothetical stage of molecular evolution, initially devoid of protein enzymes, in which all functional catalysts were ribozymes. Only one fact concerning the RNA World can be established by direct observation: if it ever existed, it ended without leaving any unambiguous trace of itself. Having left no such trace, the latest time of its demise can thus be situated in the period of emergence of the current universal system of genetic coding, a transformative innovation that provided an algorithmic procedure for reproducibly generating identical proteins from patterns in nucleic acid sequences. Today that system utilizes amino acyl-tRNA synthetase (aaRS) *protein enzymes* to attach amino acids to cognate tRNAs. However, the most extensively elaborated version of the RNA World (Koonin and Novozhilov 2009, Koonin 2011) is one in which the code was first operated by *ribozymal* aaRS, whose functions were progressively “taken over” by ancient aaRS enzymes, distant ancestors of the functional molecular species found in every contemporary living cell, mitochondria, chloroplasts and numerous viruses (Claverie and Abergel 2010, 2016; Legendre, Arslan et al., 2012).

A companion paper (Carter and Wills, 2017) outlines evidence tracing the origin of genetic coding to ancestral enzymic aaRS, and provides detailed justifications for the conclusion that such ancestry is inherently more probable, in addition to being vastly better documented, than any “takeover” by proteins of a pre-existing translation system based on ribozymes. Much of the argument rests on new comparisons of differential equations describing the translation dynamics of ribozymal, protein, and hybrid assignment catalysis and a generalization of the coupling within mathematical models for gene-replicase-translatase (GRT) systems (Füchslin and McCaskill, 2001; Markowitz et al., 2006). These elements proved too extensive to be compatible with a single publication, and are therefore developed in detail in this paper.

It is instructive to examine the process whereby a complex chemical network—i.e. ribosomal protein synthesis on mRNA templates—may have been able to survive intact while effecting a practically complete transition from one catalytic polymer (RNA) to another (protein). By undertaking such an enquiry we hope to shed light on the extent to which the major molecular biological processes of inheritance and metabolism might have been established in an RNA World, which later succumbed to the superiority of protein catalysis.

*Autocatalysis can be viewed as an essence of biology*. The complete set of molecular components needed to build a cell must be synthesized by the reaction network in the cell, starting with some basic food set available in the environment, with the second law of thermodynamics requiring that the food have higher free energy than the waste products ultimately exported back into the surroundings. The theory of the architecture of such “reflexively autocatalytic food sets” (RAF sets) has been studied in detail, starting with Kauffman (1986), and extensively elaborated in recent years by Hordijk, Steel and others [see Hordijk (2016) for a review].

The upshot of RAF theory is an impression that we should expect to find extensive, complex chemical reaction networks in nature simply as a result of the propensity of molecules, especially polymers built from a discrete set of monomers, to act as catalysts of specific chemical reactions. Indeed, cellular biochemistry seems to be rife with networked autocatalytic sets (Sousa et al. 2015). The facility with which chains of nucleotides can be copied with tolerable accuracy, along with their offering a range of catalytic (Breaker et al., 2003) and regulatory (Breaker, 2012) possibilities, lends plausibility to RNA as a substrate for RAF set formation in the prebiotic world. It is extremely hard to envisage how proteins, for which there is no known sequence copying mechanism, could form a sustainable autocatalytic network in the prebiotic world (Eigen, 1971a). Thus, life’s need to have multiple catalytic functionalities integrated into a single catalytic network has seemed, at least superficially, much easier to achieve with ribozymes than with protein enzymes.

*RAF theory provides a basic framework for describing the self-organization of reaction networks*. However, it has had little to say about the actual evolution of either biological specificity or inheritance (Hordijk et al., 2014), with similarly limited analyses of the plausible evolution of catalytic sets based on two kinds of polymers (Smith et al., 2014). Notably, none of these treatments has incorporated notions of error-prone information transfer (template replication, transcription or translation), indicating that other concepts have to be brought to bear before the advent of coded protein synthesis can be encompassed in the theory of autocatalysis. Some aspects of this problem have been discussed previously (Wills 2001; 2016), but not in relation to a possible transition from an extant RNA World. The companion paper (Carter and Wills, 2017) highlights the importance of articulating the mathematical implications of such a transition in order to properly assess the relative plausibility of different scenarios.

*On the other hand, the ideas of evolutionary coexistence and cooperation are beset by a raft of theoretical problems that frustrate attempts to explain the evolution of diverse, integrated, functional specificity in molecular genetic systems*. These problems can be regarded as chemical versions of macroscopic, population dynamic problems (group selection, kin selection, multilevel selection, inclusive fitness, altruism, parasitism, etc.) that have led to ongoing disputes between competing schools of evolutionary thought. One thing the field of population genetics has taught molecular evolutionists is that population (read “chemical concentration”) variation in spatial dimensions can be as important in determining what survives, and the internal interactions comprising the selected system’s structure, as temporal variations at a specific location in space, the latter being subject to uncomplicated rules of selection dominated by mass-action and kinetic competition (Eigen, 1971a).

As Turing (1952) first showed in relational to molecular systems, elementary reaction-diffusion coupling can result in extraordinary spatio-temporal ordering, effects which rescue some intricately cooperative chemical systems from extinction. The most extensive work on this topic has been conducted by Hogeweg and colleagues, starting with studies of hypercycles (Boerlijst and Hogeweg, 1991; 1995) and culminating most recently in treatments of parasitism (Colizzi and Hogeweg 2016a) and the evolutionary significance of high-cost cooperation (Colizzi and Hogeweg 2016b). Related studies, of paramount significance for our current undertaking, is the work of Füchslin & McCaskill (2001) and Markowitz et al. (2006) on the evolution of cell-free genetic coding in gene-replicase-translatase (GRT) systems.

*An important conceptual distinction implicit in the field of the dynamics of complex prebiotic systems is that between intrinsically and extrinsically driven self-organisation*. Darwinian selection was established as a paradigm of molecular genetic self-organisation in the pioneering work of Eigen (1971a). Differential rates of synthesis and degradation of replicating polymer variants, parameters intrinsic to the internal dynamics of the polymer population, are sufficient to establish a phase transition in the population distribution, corresponding to the survival of the fittest as a result of natural selection, conditional upon the accuracy with which the polymers are copied being above a certain threshold. However, nucleic acids (RNA and DNA) are the only polymers that are synthesized through a read-and-copy mechanism and they do not usually compete for survival directly according to their differential rates of synthesis and degradation. Rather, their survival is determined by the fitness of a much more complex replicating unit, a cell for example. A nucleic acid gene is retained if it somehow contributes to, or at least does not diminish, the fitness of the phenotype of the cells in which it is found. Under those circumstances self-organisation within the overall population of nucleic acid polymers depends on factors *extrinsic* to the process of polymer replication itself. Blind selection cannot read phenotypic properties and copy them back into genetic messages. More generally, extrinsically driven self-organisation is observed when the feedback leading to more complex individual systems (Hogeweg and Takeuchi, 2003) is externally imposed, and realized only because their internal structure happens to endow the units of reproduction, which are typically encapsulated (Szathmáry and Demeter, 1987) with a selective advantage under prevailing environmental conditions (availability of food, absence of parasites, etc.). The operational evolutionary “level of selection” (Keller, 1999; Okasha, 2006) is higher than that of any individual components. Kun et al. (2015) describe in detail the role that extrinsically driven self-organisation necessarily plays in RNA World scenarios.

In contrast, intrinsically driven self-organisation depends on interactions between components within the system encompassing the feedback that drives the system to higher complexity by amplifying some internal processes at the expense of others. Typically, the intrinsic drive to self-organisation arises out of some instability in the internal dynamics of the system: some cyclical process is tipped into a self-amplifying mode, which does not become fully damped until the system reaches a new dynamic attractor, usually a state of higher dissipation and lower entropy (Prigogine and Nicolis, 1971). The self-organised state is intrinsically stable because any incremental deviation from it temporarily produces a corresponding drive to restore it. The paradigm of intrinsically driven self-organisation of molecular phenotypes rather than genotypes is the emergence of coding in the presence of reflexive information, i.e., genes that encode a coding set of aaRS assignment catalysts (Bedian 1982; Wills 1993). The companion paper (Carter and Wills, 2017) expands our understanding of reflexivity by demonstrating that its natural occurrence is embedded in the manner in which the physical chemistry of amino acids drives protein folding to produce structural elements for the selective recognition of amino acids according to their physical properties and tRNAs depending on the presence of corresponding sequence motifs, first in the acceptor stem and later in the anticodon loop. *A genetic code operated by enzymes is differentiated decisively from one operated by ribozymes in terms of the distinction between intrinsic and extrinsic modalities of emergent complexity*.

*We begin our exposition by isolating the dynamics of codon-to-amino acid assignments from all other factors that influence the process of information transfer from the nucleotide sequences of (messenger) RNAs to the amino acid sequences of proteins*. We do not investigate how, or at what stage of prebiotic evolution relative to the development of ribosomes and metabolism, the process of collinear sequence information transfer arose. Rather, we are concerned with how coding assignments can be embedded in the chemical dynamics of a RAF system, implemented either by ribozymes or enzymes. It is quite impossible to recreate the entire path along which the very complex process of translation evolved, but it is possible to lay out certain theoretical problems that come up in trying to describe any such path, and the principles of their solution. Our approach is to extend earlier mathematical models of coding self-organization (Bedian 1982; 2001; Wills 1993; 1994; 2004) that outlined the evolution and dynamic stability of a protein-operated genetic code, presuming the existence of the extremely particular reflexive information, which expresses, according to the rules of a specified code, the specific proteins that operate that code.

We show that translation errors would inevitably be higher for any hybrid coding situation driven by both ribozymal and protein aaRS than they would be for a homogeneous system with only the initially established type of aaRS. A glossary of the nomenclature used to specify the parameters and variables of the model is provided in Table I. The main conclusion of our analysis is that the RNA World hypothesis is of little relevance to our understanding of the emergence of genetic coding as we know it, a natural feature of molecular processes that is unique to living systems. Organisms are the only products of nature known to operate systems of symbolic information processing that are essentially computational. In fact, it is difficult to envisage how alien entities found with a similar computational capability that was necessary for their existence, no matter how primitive, would fail classification as a form of “life”.

**Table I.**
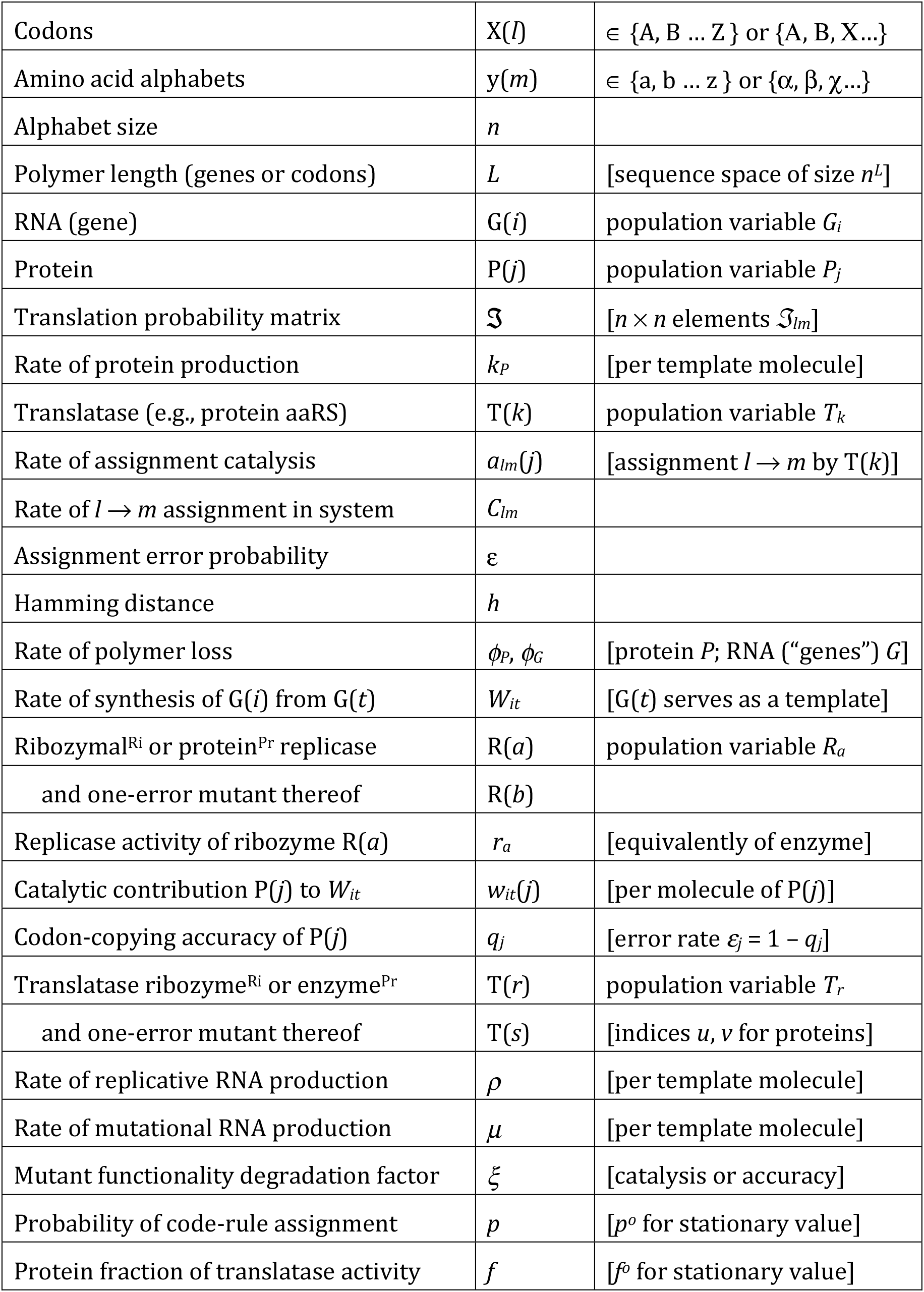
Glossary of terms

## 2 Model of translation dynamics

We consider a simplified model of protein synthesis that focuses on its information transmission aspects: translating codons at particular positions in genetic sequences into appropriate amino acids at particular positions in protein sequences. We pay particular attention to the fact that translation assignments are inherently probabilistic and are thus associated with error rates. We likewise emphasise the fact that all known amino acyl-tRNA synthetase (aaRS) enzymes are themselves proteins produced by error-prone translation of genetic sequences. We consider for simplicity a non-redundant code with one codon per amino acid. Translation assignments depend on the rates at which aaRS species, whether enzymes or hypothetical ribozymes, charge tRNAs “correctly” or “erroneously” in an inherently probabilistic fashion. Contemporary molecular biological systems approximate a true code: aaRS enzymes attach specific cognate amino acids to tRNAs bearing anticodons complementary to specific corresponding codons, with very infrequent errors.

However, collinearity of peptides with genes is a logical prerequisite for the existence of a code. It is meaningless to imagine that something corresponding to the human notion of a “code” could be instantiated in a primordial molecular biological system until such a system had acquired a reasonably stable process to maintain collinearity between an extant genetic sequence G(*i*) and the sequence P(*j*) of any peptide being synthesized. Without such a mechanism the notions of codon-to-amino acid assignments and translation have no meaning (Wills, 2016). On the other hand, evolution is not driven by logic and recent analyses of ribosomal phylogeny (Petrov and Williams, 2015; Root-Bernstein and Root-Bernstein, 2015) suggest that the required collinearity may well have emerged in a single self-organizing transition simultaneously with the most primitive form of coding. We consider in §4 a more general statement of such requirements for simultaneity in the emergence of coding.

In order to remain tractable, we ignore complications—coding redundancy; initiation and termination; variable translation rates; ribosomal stalling at rare codons or when the tRNA bearing the complementary anticodon is depleted; energetics of peptide chain elongation and ribosomal translocation, etc—that cause special effects in real molecular biological systems. We will consider only what happens when genetic sequences are translated in an implicitly synchronised step-by-step process so that proteins are synthesized at the same rate from all genetic sequences of the same length and the ribosome performs as a mechanical clockwork-ratchet device. This “clockwork ribosome” model would assure that effects we find will be due solely to the operation and stability of the translation table defining rates at which codon-to-amino acid assignments are made, whether or not they are regarded as “correct” or “erroneous”. Should it turn out that the origin of coding depends on specific features of translation and its dynamics, such as the relative sizes of different components involved in the process, then such complications will signify the need to improve the simplified model in order for it demonstrate the relevant phenomena.

### 2.1 Translation table

We define alphabets of codons {A, B … Z} and amino acids {a, b … z}, both of size n. The universal genetic code has a (sense) codon alphabet of size 61 and an amino acid alphabet of size 20, but we will start by considering alphabets of equal sizes, mostly with small values of n, especially a minimal binary code (*n* = 2). We will eventually consider the circumstances under which the number *n* of distinguishable codons and amino acids can increase or decrease.

Probabilities, 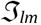, with which codon X(*l*) is translated as amino acid y(*m*) with indices *l* and *m* both running over the range 1,2…*n*, specify an *n* × *n* translation table, 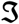. A gene G will be considered to be a sequence of *L* codons X_1_X_2_X_3_ … X_*L*_, a single point in the genetic sequence space of all *n^L^* possible genes. Translation of G produces a protein, P, whose amino acid sequence y_1_y_2_y_3_ … y_*L*_ is collinear with the codon sequence of G. Codon-to-amino acid assignments and translation acquire meaning only in the context of a reasonably stable collinearity between an extant genetic sequence G(*i*) and the sequence P(*j*) of any peptide being synthesized (Wills, 2016). Because translation is a stochastic process, repetitive translation of G produces a population distribution of proteins with different sequences.

### 2.2 Coding assignment catalysts

We assume that a ribosome produces proteins from a single genetic template at a rate *k_P_*, so that each codon on an mRNA template is translated at the same, “clockwork” rate *k_P_*/*L*. However, the rate at which any particular assignment is made depends on the relative concentrations (or population numbers) of the charged tRNAs and these depend in turn on activities *a_lm_* of all of relevant aaRS species and all amino acids present in the system. We will call the aaRS catalysts “translatases” T, whether they are ribozymes or enzymes. The rates *C_lm_* at which codon X(*l*) to amino acid y(*m*) assignments are made can be found by summing over contributions *a_lm_*(*k*) from all possible translatase species T(*k*) with individual populations *T_k_*, spanning the RNA or protein sequence space:

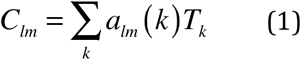

Many individual species will not be present, *T_k_* = 0, and many others will not display activity for a particular assignment, *a_lm_*(*k*) = 0. The translation table 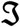 lists the probabilities 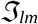 of codons X(*l*) being assigned to amino acids y(*m*) assignments and can be found by normalising the rates *C_lm_* by factors that represent the total catalytic capacity of the system for assignment of codon X(*l*) to *all* amino acids:

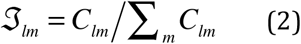

It should be noted that all of these quantities, notably the *C_lm_*, are formally functions of time, because the population variables *T_k_* are. In other words, the translation table can vary, even increase its effective dimension, on account of variations in the population of ribozymes or proteins that catalyse assignment reactions.

If each possible codon X(*l*) is translated as a corresponding, unique amino acid y(*m*), we can assign an arbitrary order for the codon indices *l* and then give each codon’s cognate amino acid the same index *m* = *l*. A population of assignment catalysts T(*k*) that executes a code in this manner consists only of ribozymes or enzymes with the property *a_lm_* = *a* ≠ 0 for *l*=*m* and 0 otherwise. Then, the table of translation probabilities is the identity matrix, 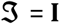:

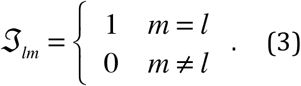

A much more realistic scenario is an imperfect coding system in which assignment errors occur at a uniform probability *ε* and the translation table has the form

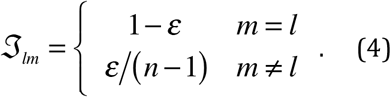

“Completely non-coding” systems make random assignments of amino acids to codons and have a translation table of the form 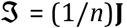, where **J** is the *n*×*n* all-ones matrix, so that

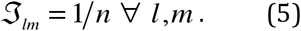

### 2.3 Code expansion occurs by phase transitions that decrease entropy in sequence space

Coding self-organization and code expansion can occur in systems whose translation dynamics have particular instabilities, across which a transition can occur from one attractor state to another in which the translation rate matrix has higher effective dimension, *n_eff_*. Such transitions increase coding capacity, hence the precision with which any set of genetic messages is translated, thereby facilitating eventual reductions in the sequence space occupation—and entropy—of the corresponding ribozome or proteome. As an example, in the non-coding reference system all *n* codons are indistinguishable, so the effective number of codons is *n_eff_* = 1. In these terms, transition from a noncoding dynamic state to the operation of a binary code (*n*=2) is a special case of single-step code expansion: the effective dimension of 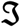 increases by one. Delarue (2007) discusses code expansion broadly, in terms consistent with an initial binary differentiation of amino acids according to their attachment to cognate tRNAs by separate Class I and II aaRS enzymes.

The possibility of code expansion transitions has been demonstrated in several computational models whose dynamics are contingent upon protein-based aaRS translation, with a variety of rules describing how the catalytic capabilities *a_lm_*(*j*) of a protein P(*j*) depend on its amino acid sequence (and implicit folding). The first study (Wills, 1993) concerned systems whose genetic information was fixed. Three different “embeddings” for the association of *a_lm_* values with points *j* in the protein sequence space were considered: (I) random; (II) constant activity *a* within a fixed Hamming distance radius *h*_0_ around *n*^2^ randomly chosen centres, one for each possible codon-to-amino acid (*l* → *m*) assignment; and (III) also with randomly chosen centres of *l* → *m* assignment activity, but with *a* defined as a narrow Gaussian function exp 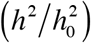, rapidly decreasing beyond Hamming distance *h*_0_ from the centre but non-zero even at the furthest extremity of sequence space. System dynamics in each case were demonstrated to have non-coding and coding attractors separated by an instability. In all cases the genetic information supplied to the simulated system was constructed so that it was reflexive for a code that was chosen in advance.

Reflexivity means that when the genetic information is translated according to the rules of some code the protein products’ joint activities constitute a translation table for that specific code, as in Eq (3) or (4). In relation to case (III) above, it was further demonstrated (Wills, 1994) that progressive loss of strict reflexivity due to mutation of the genetic information leads to a progressive increase in the coding error rate *ε*. And a separate study Wills, (2004) investigated systems in which nested instabilities allowed for a series of code-expanding transitions to attractor states with progressively larger values of *n_eff_*. That study focused on a further case, (IV): as a result of spontaneous changes in the translatase populations (transitions described as “quasi-species bifurcations” in the companion paper: Carter and Wills, 2017), each letter of binary alphabets of codons {A, B} and amino acids {a,b} can bifurcate into two versions to produce four-member codon and amino acid alphabets, {A, B, X, Δ} and {α, β, χ, δ}, increasing the coding capacity from 1 bit to 2 bits, and expanding the 2 × 2 translation table into a 4 × 4 table. The hierarchically nested embedding of assignment activities used for that simulation geometrically mirrored the decomposition of the alphabets, showing stepwise coding selforganization, first from a non-coding state to the execution of a binary code {A→·a, B→·b} and then from the binary code to the expanded four-dimensional code {A→α, B→β, X→χ, Δ→δ}.

### 2.4 Differential equations for translation dynamics

The rate at which the population (or concentration) *P_j_* of a protein with the particular sequence P(*j*) = y_1_y_2_y_3_ … y_*L*_ changes as a result of being synthesized through translation of a single stranded nucleic acid template with sequence G(*t*) = X_1_X_2_X_3_ … X_*L*_, and consequently lost (the *φ_p_* term), is given by

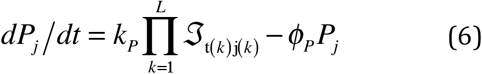

where *l* = t(*k*) is the identity of X_*k*_, the codon that occurs at position *k* in the gene G(*t*), *m* = j(*k*) is the identity of y_*k*_, the amino acid that occurs at position *k* in the protein P(*j*), and *φ_p_* is the overall rate of loss of P(*j*) due to degradation and movement out of the system, considered constant across all species. For a perfectly operating code, the net rate of production of every coded protein is simply *k_p_* − *φ_p_P_j_* per encoding gene but as noted above, this situation exists only when the aaRS catalysts for the code have perfect specificity and none of the genes encodes a catalyst for an assignment function not in the coding set.

In the case of an imperfectly operating code we must sum the effects over all gene templates G(*t*) present in the system because, with errors, there is a non-zero probability, no matter how small, of any protein resulting from the translation of any gene:

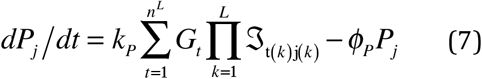

Substitution into Eq (6) of the expression for 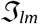 from Eqs (1–2) yields a system of nonlinear differential equations for protein production, which is the basis of coding self-organisation in an evolving PCW (Wills 1993): in the presence of a fixed set of genes displaying a significant degree of reflexivity, the aaRS assignment catalysts that the genes (at least partially) encode embody an autocatalytic set whose population is self-amplifying beyond the threshold of stability— stochastically accessible by local fluctuations in 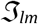—separating basins of attraction of non-coding and coding dynamic states (Wills, 1993). In any case, the solution of Eq (7) corresponding to the coding state describes the maintenance of the suite of proteinaceous aaRS assignment catalysts {T^Pr^} that operate the genetic code in living cells.

### 2.5 Coupling replication to translation: the need to copy information

Any realistic representation of molecular turnover under prebiotic conditions must take into account the error-prone replication of genetic information and its accumulation and preservation as an outcome of natural selection. Thus, Eq (7) must be supplemented by a similar system of equations describing the net rates of increase of gene sequences.

When genes are replicated, their population dynamics may be described by the quasispecies equation (Eigen 1971a), which is in many respects similar to (7):

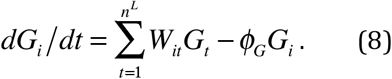

The coefficient *W_it_* is the rate at which G(*i*) is synthesized when G(*t*) serves as a template for the replication of a nucleic acid molecule of length *L*. When nucleic acid replication is accomplished by a process that produces new sequences from each template at a rate *r* with codon-copying accuracy of *q* (or error probability *ε* = 1 − *q*) the coefficients *W_it_* can be specified in terms of Hamming distance *h_it_*, measured in terms of codon differences, between G(*i*) and the template G(*t*). Assuming random errors in codon copying we have

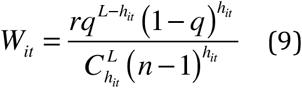

which is a modified form of Eq (27) of (Eigen et al., 1988). Eq (8), like Eq (7), represents a nonlinear system of differential equations, in this case describing Darwinian competition between sequences. The steady state solution of Eq (8) circumscribes a population of polymer sequences comprising a quasispecies, a population distribution comprising a “cloud” in sequence space centered on a consensus sequence defined by its relatively optimal functionality. Whether or not that centroid sequence is the dominant sequence, depends on the magnitude of the error rate *ε*. Above a certain threshold in *ε* the steady state population degenerates to a uniform distribution across the entire polymer sequence space. That threshold has been called Eigen’s Cliff’ (Koonin, 2011). It is a form of ‘error catastrophe’ considered in a wider range of circumstances (Orgel, 1963).

If nucleic acid copying is accomplished by a replicase, then in an RNA World any such catalysts must themselves of necessity be nucleic acids. Let us designate a subset of nucleic acid sequences R(*a*) that can act as general replicases, such as are being investigated experimentally with increasing success (Horning and Joyce 2016). Eq (9) is now rewritten in terms of specific replicase activities *r_a_* and codon-copying accuracies *q_a_* which define rate coefficients *w_it_*(*a*) appropriate to each replicase R(*a*), template *G_t_* and RNA gene product *G_i_*. The coefficients in Eq (8) then become

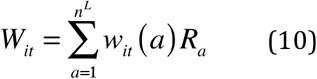

Substitution of Eq (10) into (8) gives

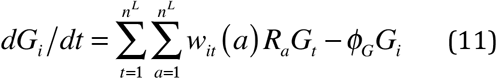

which is the standard “replicator equation” affording the most general description of the basic dynamics of an autocatalytic network of ribozymes. It says that any RNA molecule of length *n* codon units can serve as a template G(*t*) for the production of any other RNA molecule G(*i*), depending on the mutation rate characteristic of each ribozymal replicase R(*a*) that replicates the template in an error-prone fashion.

## 3 Evolutionary dynamics of ribozyme-dominated systems

Takeuchi and Hogeweg (2012) provide an excellent analysis of the evolutionary dynamics of systems of “replicator equations” like Eq (11), including hypercyclic interdependence (Eigen 1971a, Eigen and Schuster, 1979) as well as mechanisms of cooperation that must operate to overcome the effects of Darwinian competition in molecular systems, especially in an RNA World. It is first necessary to take into account the discrete character of population variables such as *G_i_*, which are treated as continuous variables in ordinary differential equations. The ODE approach fails to represent important discontinuous phenomena like extinctions, especially in small, localized populations. With that proviso their conclusions are salutary:

i. Even with artificially imposed hypercyclic feedback, global maintenance of mutually dependent molecular species cannot be expected. Non-functional “parasites” are unavoidably replicated in the system and exacerbate this problem, as they contribute insufficiently to replication to compensate for the necessary energetic investment.
ii. Parasitism can be overcome by spatial differentiation to create systems of interdependent replicators and a higher level at which Darwinian selection operates, the system level rather than that of individual molecules. When the individual population variables *G_i_* depend on spatial variables (*x*, *y*, *z*) on a scale comparable to distances that molecules traverse during single molecular “birth-death” processes, systems of cooperating replicators can survive as spatially self-organized, mesoscopic waves, with various geometries (Turing, 1952), that continually “purify” themselves of parasites.
iii. Parasites nevertheless furnish mutational bridges for novel replicator species to enter into the system.
iv. If molecular reactions are spatially compartmentalized, higher level competition determines selection explicitly for survival between replicating compartments rather than individual molecular species. Compartmentalization can thus both solve the problem of parasites and provide for the “division of labour” between templates and catalysts, and the emergent separation of these roles into different polymer types.

### 3.1 Proteins in an RNA World

It is generally assumed that the indirect effect of improving metabolic catalysis achieved by the introduction of enzymes into an RNA World in which everything was previously accomplished by ribozymes was sufficient to give the requisite advantage to protein-encoding genes in the competition for survival among competing RNAs. The manner in which proteins, produced according to Eq (7), could have such an influence in an RNA World is represented by the constants *w_it_*(*a*) in Eq (11), the rate at which a replicase R(*a*) produces the RNA species G(*i*) using G(*t*) as a template. That is to say, the proteins must exert their effect by influencing the outcome of RNA replication events in which they serve no necessary role. For the time being we will set aside the evident problem of how protein parasites could self-organise to coerce an RNA world into inventing a system for their coded synthesis, including the accumulation and preservation of a body of RNA information in which they were encoded, solely through their effect on the *w_it_*(*a*) parameters.

In order for an “RNA Coding World” to exist, prior to the evolution of proteinaceous aaRS enzymes, various individual RNA species within the surviving population must serve the roles of aaRS assignment catalysts (“translatases”) and genetic templates for the production of proteins. Substitution of Eq (2) into (7) gives

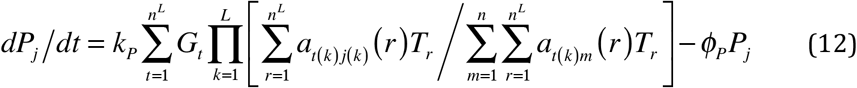

assuming that a suite of ribozymal translatases T(*r*) specifically attach amino acids *m* to tRNA-like adaptors bearing anticodons to *l* at characteristic rates *a_lm_*(*r*). However, the full system of dynamic equations (Eq 12) describing this scenario is intractable without additional assumptions. Therefore, let us adopt a heuristic approach similar to that of Takeuchi and Hogeweg (2012) and consider a much simplified case.

Suppose a population of ribozymal translatases makes “correct” codon-to-amino acid assignments with overall probability *p* = 1 – *ε*, according to the rules of a chosen code. Then, the translation matrix for the system will be of the form

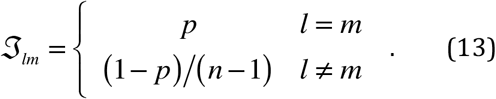

which is the same as Eq 4. The rate of production of any protein P(*j*), given the presence of a gene G(*j*) encoding it and neglecting mutants of G(*j*), will then be given by a much reduced form of Eq (12):

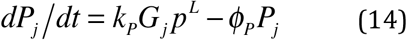

The convex surface defined by (14) has a stationary solution at *P_j_* = *G_j_* (*k_p_ p^L^*/*φ_p_*) that is stable, showing that the production of proteins in an RCW depends on the system’s ability to maintain, in addition to functional ribosome-like machinery, not only the genes encoding the proteins but also accurate ribozymal aaRS catalysts. The translatase T(*r*) and the protein encoding genes G(*j*) must, through some trick of self-organisation, overcome their role as parasites in a system of replicators that has to compete with *all* species that it replicates.

Survival evidently requires the operation of some mechanism of self-organization that selectively keeps local populations of the parasitic species low enough for the replicators to sequester adequate resources for their own reproduction (Takeuchi and Hogeweg 2012).

### 3.2 Ribozymal replicases (replicators)

Consider a simple RCW: a perfect ribozymal replicase R(*a*) that makes no copying errors maintains a population of ribozymal translatases T(*r*) that form a coding set, providing for the synthesis of proteins that are encoded in genes G(*i*), “genetic information” that has been selected on account of the advantage conferred on the system by the proteins’ contribution to its functioning and maintenance. Eqs (8 & 11) for the time evolution of the ribozymes reduce to

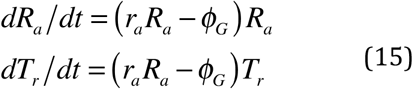

where *r_a_* is the rate at which a single replicase molecule copies a single copy of a single gene. The only significant dynamic fixed point of the system exists at *R_a_* = *φ*_G_/*r_a_* whose stability depends on how the loss rate *φ_G_* is constrained. While *R_a_* is regulated, the value of *T_r_* will be subject to stochastic drift, leading eventually to extinction of the translatases due to the usual finite population size effects (Küppers, 1983). However, the same argument applies to any finite population of perfect replicators, even if there is more than one such, G(*a_i_*: *i*=1,2…), so R(*a*) will meet the same fate, unless some mechanism of self-organization allows the system to gain advantage from cooperation between the different polymer types (Hickinbotham and Hogeweg, 2016). We continue by assuming that is the case.

If we suppose that the replicase R(*a*) copies RNA strings with an accuracy of *q_a_* = (1 − *ε_a_*) per codon and that the replicase functionality of one error mutants R(*b*), is reduced to a fraction *ξ_a_* of that of R(*a*), in respect of both catalytic capability and accuracy such that *r_b_* = *ξ_a_r_a_* and *q_b_* = *ξ_a_q_a_*, then the time evolution of the replicase ribozymes is given by

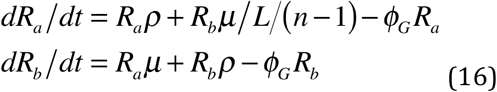

where the rates of replicative and mutational production, *ρ* and *μ*, respectively, are given by

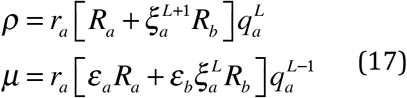

and we have taken account of single error events, including back-mutation from R(*b*) to R(*a*), but neglected all multiple-error replication processes and their effects. These equations have a fixed point (*dR_a_*/*dt* = *dR_b_*/*dt* = 0) at 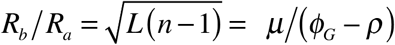. The overall probability of correctly copying codons (X) is given by

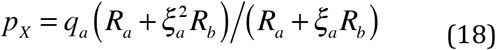

For either *ξ_a_* = 1 [R(*b*) identical to R(*a*)] or *ξ_a_* = 0 [R(*b*) inactive], *p_X_* = *q_a_* as expected; otherwise the minimum stationary value of *p_X_* occurs according to the condition 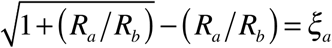.

### 3.3 Ribozymal translatases

Similarly, the rates of production of a ribozymal translatase T(*r*) and its one error variants T(*s*) in an RCW are given by Eq (16) with *T_r_* replacing *R_a_* and *T_s_* replacing *R_b_* and *ρ* and *μ* still defined in Eq (17) in terms of *R_a_* and *R_b_*. Unsurprisingly a translatase population ratio of 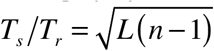 is stationary. The same conclusion applies to the population ratio for any gene relative to that of its one-error mutants when they are maintained by replicases R(*a*) and R(*b*). This applies to genetic templates G(*t*) that continue to survive in an RCW on account of the contribution that their encoded proteins makes to the viability of the selected unit of which they are a part.

If the ribozymal translatases T(*r*) and T(*s*) catalyze codon-to-amino acid assignments (X → y) with characteristic rates *a_r_* and *a_s_* = *ξ_r_a_r_* and accuracies *q_r_* = (1 − *ε_r_*) and *q_s_* = *ξ_r_q_r_* = (1 − *ε_s_*), then the translation table is of the form

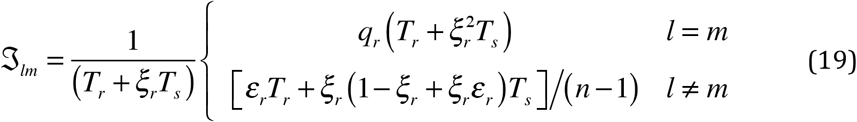

and the overall probability of making a correct codon-to-amino acid assignment is

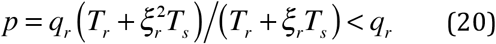

which, as expected, resembles Eq 18 very closely.

Equations (15–20) clearly demonstrate the manner in which the dynamics of an RNA world, illustrated in Fig 1(a), is completely locked into the fate of the competing replicators R(*r*,*s*…). The functional accuracy *p* of assigning the “correct” amino acid is enslaved to the ratio *T_s_*/*T_r_*, which is, in turn, enslaved to the ratio *R_b_*/*R_a_* that characterizes the functionality of the replicase population.

**Figure 1.**
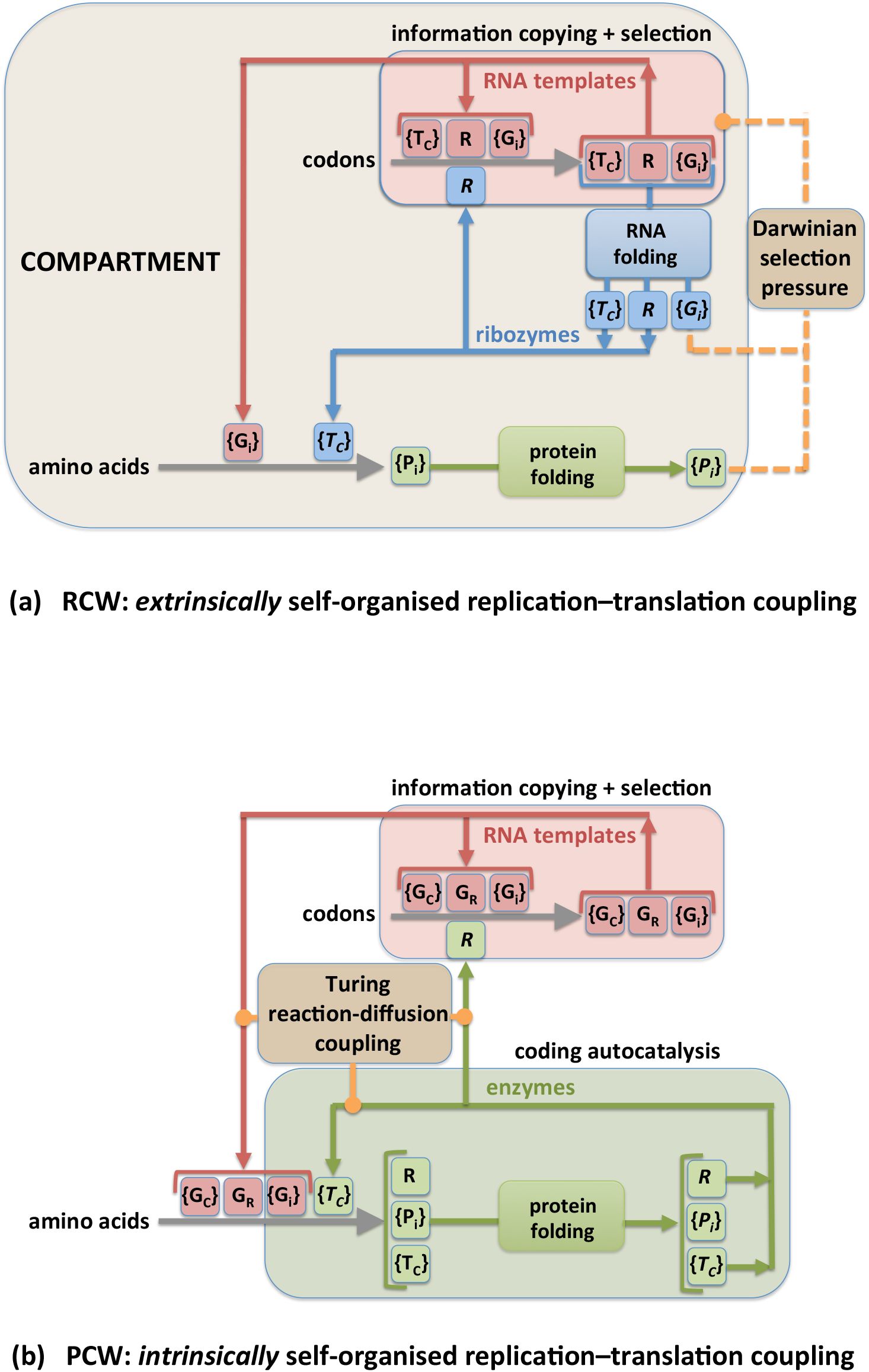
Dynamical architecture of encoded protein production. (a) In the hypothetical RNA Coding World (RCW), RNA templates (red) are copied by a ribozymal replicase R, or network of ribozymes (blue), all of which are copied by the same catalytic process. Some templates {G_i_} serve as genetic information for the production of proteins {P_i_} with amino acid sequences collinear with the {G_i_}, through a process utilizing a suite of aaRS-like translatase ribozymes {*T_c_*} that execute the rules of a code. Both RNA template sequences and protein sequences acquire funtionality as a result of spontaneous folding processes. Information processing (storage, copying, assignment catalysis) emanates exclusively from the RNA domain, with which proteins are coupled (yellow) only through their potential to exert some differential selective pressure during template copying. The origin and stability of the organised network necessarily relies on its being encapsulated and outcompeting variant forms under *extrinsic* environmental pressure. (b) In the Protein Coding World (PCW) of cellular molecular biology, template copying is catalysed by a protein (green) enzyme replicase *R* or suite thereof, for which a (red) template G_R_ carrying specific genetic information must be preserved. The aaRS translatase enzymes {*T_c_*} are also proteins requiring the preservation of templates {G_*c*_} in which their amino acid sequences are genetically encoded. In the PCW, the storage, copying and translation of genetic information all require the coupling of the RNA and protein domains. Studies of GRT systems have demonstrated that reaction networks with this information-processing dynamical architecture have an *intrinsic* tendency to self-organise into stable, spatially differentiated structures as a result of the inexorable internal operation of the Turing reaction-diffusion mechanism.

We observe that the stationary condition 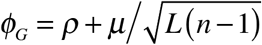 cannot be satisfied under the constraint that the rate of depletion of the products from any template be equal to the overall rate of production from that template, that is *φ_G_* = *ρ* + *μ*, except when *φ_G_*, *ρ* and *μ* are all zero, the point of extinction, or when unconstrained growth reaches its infinite limit. This illustrates the absolute need of an RCW for some form of extrinsic self-organisation to overcome the extinction problem (Hickinbotham and Hogeweg, 2016) and allow R(*a*) to overcome the problem of R(*b*) parasitism so that stable, finite values of *R_a_* and *R_b_* can be achieved, whereupon *T_s_*/*T_r_* and *p* will be constrained by the ratio *R_b_*/*R_a_*.

## 4 Dynamics of enzyme based systems

### 4.1 Imperfect protein translatases

Consider now the alternative situation in which the aaRS assignment catalysts are protein enzymes T^Pr^(*u*) rather than ribozymes T^Ri^(*r*). Obviously, the maintenance of the genes G(*u*) encoding these translatases will suffer the same problem of parasitism as they do in an RCW that uses ribozymal translatases. However, there is an additional problem. Because these translatases are proteins, their production is susceptible to an error catastrophe in the accuracy of translation, even when the exact information encoding them is not threatened by copying errors during replication. This problem is postponed until §5.

We will take account of first order errors in translation by considering one-error variants T(*v*) of the “wild-type” aaRS T(*u*) that perform the same function as T(*u*) but with activity *a_v_* = *ξ_u_a_u_* and accuracy *q_v_* = *ξ_u_q_u_* both degraded by a factor *ξ_u_* < 1. Under these circumstances the translation table 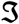 has elements given by Eq (19) with *T_u_* and *T_v_* substituting for *T_r_* and *T_s_*, respectively, and the same substitutions into Eq (20) give the expression for the probability of correct coding assignments, *p* < *q_u_*.

The expressions for the production of the translatase proteins are obtained from Eq. (7) as

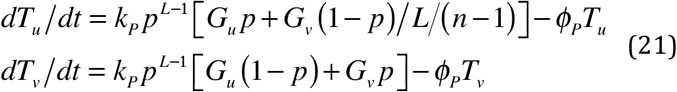

where the second term in the expression for *dT_u_*/*dt* takes account of the rare events in which mistranslation of a mutated gene results in production of a functional protein. In order to find the fixed points of this system we set *dT_u_*/*dt* = *dT_v_*/*dt* = 0, solve for *T_u_* and *T_v_* in terms of *p* and substitute these into Eq (20) to obtain the polynomial

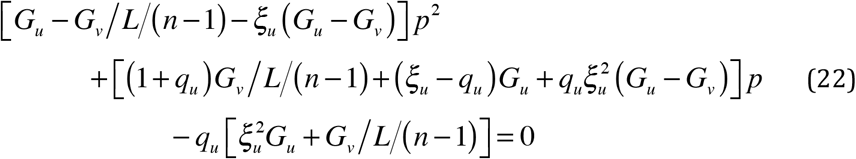

The root of Eq (22) corresponding to the physically realistic steady state (“*o*” superscript) is in the range *ξ_u_q_u_* ≤ *p^o^* ≤ *q_u_*. For *ξ_u_* = 1 (one error mutants indistinguishable from wild-type) and *ξ_u_* = 0 (one error mutants inactive), it occurs at *p^o^* = *q_u_* as expected. The stability of the fixed point requires *∂*(*dp*/*dt*)/*∂p* < 0, a condition which can be explored through appropriate manipulation of Eqs (20–22) to show that the required inequality simplifies to *p* > *p^o^*(*L* − 1)/*L*, which is clearly satisfied by *p* = *p^o^*, so the fixed point is stable.

### 4.2 Mixed ribozymal and enzymatic assignment catalysis

We now consider a mixed system in which there are both ribozymal T^Ri^(*r*) and protein T^Pr^(*u*) translatases which effect codon-to-amino acid assignments at different characteristic rates *a_r_* and *a_u_*, respectively, and accuracies *q_r_* and *q_u_*, respectively. Note that the populations of T^Ri^(*r*) and T^Pr^(*u*) are required to operate the same set of *n* codon-to-amino acid assignments and T^Pr^(*u*) is critically dependent on the genes G(*u*) being maintained in the RNA population. We will neglect the effects of ribozymal mutants T^Ri^(*s*) and protein variants T^Pr^(*v*). The form of Eq (19) relevant to mixed ribozymal-enzymatic translation is

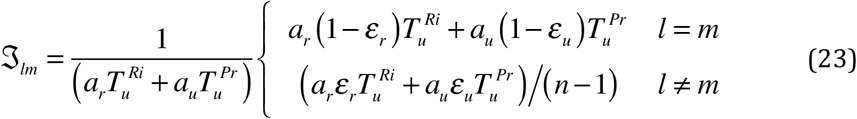

and the accuracy of coded translation is

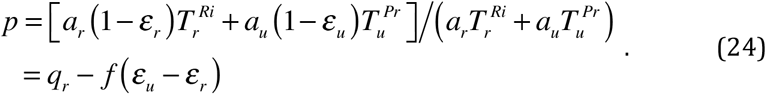

where 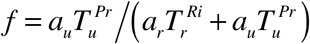 is the fraction of codon-to-amino acid assignments that are effected by protein T^Pr^ rather than ribozymal T^Ri^ catalysts. The time evolution of the ribozymal and protein translatases is given by

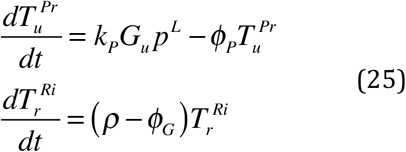

and the dynamic fixed point of the translation process corresponds to the solution of the polynomial equation

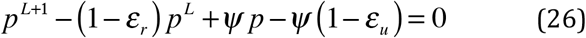

that lies in the range *q_u_* < *p* < *q_r_* (presuming *q_u_* < *q_r_*). Eq (26) is obtained from Eqs. (23) and (25) and the coefficient of the first order term is 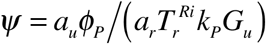). The steady state accuracy of translation can also be expressed implicitly in terms of 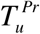 as

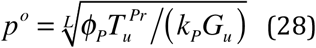

which is the *L*^th^ root of the ratio of the rate of loss of the protein translatase to the rate of production of proteins using the translatase-encoding gene G(*u*) as a template-message for translation.

These results, especially Eq (24), demonstrate the degrading influence of a less accurate protein coding system on a coexistent ribozymal system. The ribozymal coding system will function autonomously, dependent only on maintenance of the ribozymal translatase species T^Ri^(*r*). However, the protein coding system T^Pr^(*u*) regulates the overall accuracy of coding. The dynamic system has an attractor state in which the protein population makes a contribution *f^o^k_p_* to the overall rate of translation of any genetic template and a contribution of magnitude *f^o^*(*ε_u_* − *ε_r_*) to the overall error rate 1 − *p^o^*. In other words, whatever selective advantage coded proteins confer on a system incorporating a ribozymally operated system of translation, the inception of less accurate enzymic translatases will diminish that advantage, favouring variant systems that lack both the proteinaceous aaRSs and energetic load involved in maintaining the genes that encode them.

Finally, it should be noted once again that any protein coding system is dependent on the maintenance of a population of templates G(*u*) that encode the protein translatases T^Pr^(*u*) and that both the ribozymal species T^Ri^(*r*) and the translatase-encoding species G(*u*) are required somehow to survive, essentially as parasites, in a world of RNAs replicators. It is the dynamics of gene replication and its effect on the evolution of translation to which we now turn our attention.

### 4.3 Mixed ribozymal and enzymatic protein replicases

The question of genetic information copying is at the heart of the Darwinian evolution. Let us briefly consider the advent of a protein replicase R^Pr^ in a functional RCW in which relatively sophisticated and accurate information copying has evolved through selection of a general ribozymal replicase R^Ri^, whose catalytic efficiency parameters *r_R_* and codon-copying accuracy parameters *q_R_*, produce a high probability, 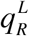, of correctly copying a genetic template. Introducing a protein replicase R^Pr^ with parameters *r_p_* and *q_p_*, the overall production of any genetic sequence G(*i*) will, neglecting back-mutation processes, be (c.f. Eq 11)

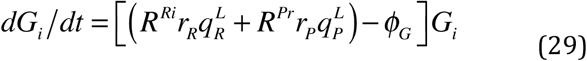

and the probability of correct copying will be

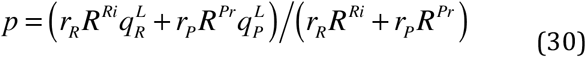

The problem here is the same as with the advent of protein translatases [Eq (24)]: as long as *q_p_* is significantly less than *q_R_* which is to be expected for the first proteins showing replicase activity to emerge in an RCW, the probability of correct copying will be diminished for *all* genes, but especially the ribozymal replicase R^Ri^. Since the evolution of the system has been optimized under the constraint of the variables *r_R_*, *q_R_* and *R^Ri^*, all of which will be expected to diminish, the system will be at risk of sudden catastrophic disintegration unless it quickly purges itself of the emergent protein replicase by purifying selection.

## 5 The self-organizing RNA-Protein world: GRT Systems

The discussion in §2 recapitulated the requirement established in studies of coding self-organisation (Bedian, 1982; Wills, 1993) for feedback-constrained bootstrapping in relation to the stability of genetic coding as it occurs in living cells. Likewise, §4 developed the conclusion that it is virtually impossible for two discrete molecular genetic systems to couple constructively if they process information with substantially different error rates. In this section, we recognize that the two arguments are intrinsically complementary requirements for efficient self-organization between genetic information and the utilization of proteins for genetic coding. If we consider that errors in information transmission play a role analogous to that of impedance in relation to self-organization, we see that, as the power transfer in dissipative electronic structures is optimal when input and output impedances match, the molecular biological organization observed in living systems would have proceeded more efficiently and hence more rapidly, if at all stages of development, the effects of information processing errors in the replication of nucleic acids matched those in protein synthesis. This leads to the conclusion that genetic coding and information copying using functional proteins are most unlikely to have emerged from a world in which these functions were already performed at a higher level by ribozymes. The increase in the impedance of information transmission caused by the advent of either primitive translatase or replicase enzymes would have destabilized whatever transmission efficiency had been achieved in an extant ribozymal system. In any case, the RCW does not have the coupling necessary to establish information impedance matching between the separate processes that replicate and translate genetic information.

Living systems produce proteins P(*i*) from information encoded in genes G(*i*) using aaRS enzyme translatase T^Pr^(*u*) and their nucleic acid genes are copied using protein enzyme transcriptases and replicases R^Pr^. The dynamics of such systems can, in terms of the simplifications of the current analysis, be represented by the equations below (LHS: 31a, 32a), which are compared and contrasted with equations describing the equivalent processes in an RCW (RHS: 31b, 32b). The dynamics of the RNA domain of the PCW and RCW are described by

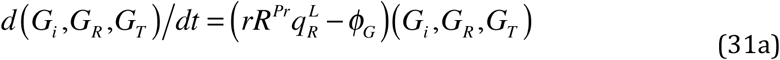

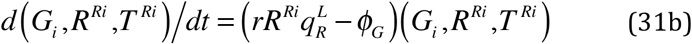

and the dynamics of the protein domain by

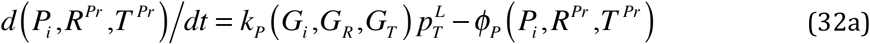

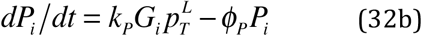

The processes described by these equations are illustrated in Figure 1. It is evident that the LHS equations for gene and protein production in the PCW are tightly coupled through the population variables *G_R_*, *G_T_*, and *R^pr^* which together represent the genes encoding R and T and the replicase enzyme itself; and that the translatase part of the protein domain is autocatalytic (through *p_T_*). However, in the RCW the dynamics of the RNA activity—replication of genetic information and catalysis of coding assignments—is completely autonomous; events in the protein domain have no effect on the value of any variable in Eq (31b). Furthermore, the protein domain is completely dependent on the RNA domain through the variables *G¡* and *pτ*, which represent the populations of encoding genes and the accuracy of the ribozymal translatase population, respectively. It is hard to envisage how, considering the elaborate feedback elements furnished uniquely in the PCW, the slow and blunt instrument of selection pressure that proteins might exert on the fitness of nucleic acid sequences in an RNA World could mold a refined system of genetic coding [cf. Fig 4 of the companion paper, (Carter and Wills, 2017)].

On the other hand, the double risk of an error catastrophe, especially at high error rates, either in information storage (replication) or encoding (translation), or both simultaneously, appears to remain a significant barrier to achieving dynamic stability of the highly coupled PCW of molecular biology, described by Eqs (31a) and (32a). What kept the emerging world of cellular molecular biology stable when the processes needed to produce the biochemical systems that operate in living cells were so precarious?

The answer to this question is to be found in territory that is unfamiliar to most biologists: the thermodynamics of highly dissipative systems, i.e., metabolically driven systems. The biological relevance of this way of thinking was highlighted in two expositions that appeared back-to-back 45 years ago, the first (Prigogine and Nicolis, 1971) outlining how order can spontaneously emerge from disorder in thermodynamically driven systems with highly coupled dynamics (like Eqs 31a and 32a) and the second (Eigen, 1971b) demonstrating that beyond a certain threshold of digit-copying accuracy, Darwinian selection among replicating polymers drives a system into a dynamic state in which the surviving quasi-species protects a certain quantum of polymer sequence information from degradation due to mutation. The point of the latter observation is that polymer replication is a metabolically driven dissipative process in which there is continual turnover of individual sequences.

Under those circumstances natural selection acts as a self-organizing force that first facilitates takeover of the system by the dominant quasi-species, i.e., the “fittest” individual and its mutant cloud, and second, staves off the error catastrophe that would otherwise threaten genetic information storage (Eigen, 1971a). When this dual principle is transferred from the domain of polymer sequence replication (Eqs 31a and 31b) to the domain of translation in the PCW (Eqs 32a), coding self-organization (Wills, 1993) can be identified as the force that enables a set of code-executing enzymes (i) to take over protein production, and (ii) to stave off the translation error catastrophe that would otherwise lead to the demise of the entire process of protein synthesis (Orgel, 1963; Wills, 1994). The idea that natural selection explains both (i) the accumulation of genetic information, and (ii) its maintenance, has been consolidated in scientific thinking over a period of more than a century and a half. However, the origin and stability of translation have seldom been regarded as a dynamic problem beyond recognition that there is a chicken-egg paradox to be resolved (Rodin and Rodin, 2008), even though the protein synthetic apparatus evidently forms a highly connected autocatalytic network likely to have diverse modes of stable and unstable behaviour.

The tightly coupled nonlinearities inherent in the equations describing the maintenance of both ordered nucleic acid and protein sequences in the world of real molecular biology (Eqs 31a and 32a) strongly suggest that there is an inherent organization among the dynamic forces responsible for the stability of the process. What is most significant in that regard is the possibility of a dire collapse: errors in translation are likely to produce replicases that function erroneously and lead to mutations in translatase genes, coupling the Orgel and Eigen error catastrophes in a vicious circle. Indeed, as Füchslin and McCaskill (2001) noted in their study of gene-replicase-translatase (GRT) systems, the only physically reasonable solution of Eqs 31a and 32a, as they stand, is the attractor state at the origin, where all species have gone extinct. However, in fulfillment of the expectation that these equations are capable of describing a generative as well as a degenerative dynamic pathway, these authors demonstrated that one need only recognize that all of the concentration variables are functions of spatial coordinates (*x*, *y*, *z*) and that molecular Brownian motion demands the inclusion of diffusion terms in all of the chemical rate equations; and then, there *are* solutions of Eqs. 31a & 32a corresponding to the stable maintenance of not only (i) the reflexive genetic information necessary for the operation of encoded protein synthesis but also (ii) the gene which encodes the replicase, the species most directly involved in arbitrating the reproductive fitness of genetic information generally. In other words, the dynamic coupling in GRT systems is so strong that the elementary physical connection between molecular turnover (chemical reaction) and movement (diffusion) is adequate to cause self-organisation of the coupled computational processes of replicating nucleic acid polymer sequence information and translating it to synthesize functional proteins, the quintessence of molecular biology.

The GRT dynamical architecture of the molecular biological groundplan is intrinsically self-organizing. Unlike the hypothetical RCW it does not require special extrinsic arrangements at some higher level of selection to assure the survival of diverse genes and functional components in compartments that replicate as complete units. Reaction-diffusion coupling (Turing, 1952) both generates and maintains the spatially variegated association of different components of PCW-like GRT systems that leads to their cooperative functionalities and protects them against the destructive effects of parasitic variant molecular forms. However, the particular efficacy of the Turing mechanism for GRT systems is grounded in an even stronger dynamic coupling between gene replication and translation. A stationary replicase population supports exponential order growth of gene populations and a stationary gene population supports exponential order growth of translatase enzymes. But when genetic selection improves coding, thereby increasing the exponential rate constant for translatase autocatalysis, and coding self-organisation supports production of an improved replicase, thereby increasing the exponential rate constant for replication, the entire system can display hyperbolic order growth due to cyclical feedback in the relication-translation meta-loop in a manner directly equivalent to that seen in hypercycles (Eigen, 1971a).

Thus, the dynamics of the PCW entail a direct, very rapid, intrinsically generative pathway to a self-organised state of encoded information processing, whereas there is no such possibility inherent in the dynamics of the RCW. And it is worth mentioning that bi-directional coding of Class I and II aaRS enzymes (Rodin and Rodin, 2008) provides a natural mechanism for the elimination of problematic competition between the two translatase genes needed for an initial binary code. On the other hand, these factors do not plausibly delineate a continuous pathway directly to anything as sophisticated as the universal genetic code. Various stages of the emergence of the code must have been associated with equivalent stages in the evolution of metabolic pathways and their control. Transitions between coding states of increasing complexity may have been driven by kinetic processes of hyperbolic order, but these would have been “punctuations” between “equilibria” in coding evolution, during which natural selection acted as the main mechanism for the adaptation of other processes to the new possibilities of computational control. Under those circumstances neither replication nor translation could be expected to maintain stability unless each helped limit the impacts of the other’s error rate, that is, unless the coupling of the flow of information between replication and translation was “impedance matched” (Carter and Wills, 2017).

## Concluding remarks

The direct evolution of a coupled world of genetic information and encoded functional proteins in real-world molecular biology is far more plausible than any scenario in which there was an initial RNA World of ribozymes sophisticated enough to operate a genetic code. The preservation of encoded information processing during the historically necessary transition of any such system to the ancestral aaRS enzymes of molecular biology appears to be impossible, rendering the notion of an RNA Coding World scientifically superfluous. While this conclusion is grounded in an understanding of exactly how the dynamical architecture of molecular biology can solve the computational chicken-egg paradox of code evolution, it leaves a host of problems concerning the evolution of the complex apparatus of translation unresolved. On the other hand, recognition of the role of reflexivity in driving the intrinsic self-organisation of molecular biological coding has stimulated a deeper enquiry into the relationships between structural determinants of the aaRS coding apparatus (Carter and Wills, 2017). The formal requirement for reflexive information that encodes assignment catalysts according to the rules of the code they execute is grounded in a more elementary, physical reflexivity. Instantiation of the computational requirement of reflexivity in the dynamic processes of real-world molecular interactions demanded of nature that it fall upon, or we might say “discover”, a self-amplifying set of nanoscopic “rules” for the construction of the pattern that we recognize as “coding relationships” between the sequences of two types of macromolecular polymers. However, nature is innately oblivious to such abstractions: the matching of amino acids to codons is achieved by folded aaRS structures that are, at least according to quantum mechanical demands, “accidentally” produced through the computationally controlled placement of amino acids with different physical properties in specific positions of variants of two basic protein folds, labeled “Class I” and “Class II”. Even this simplest of distinctions had to be a discovery of itself, a “bootblock” that could be built upon and elaborated into the improbably refined system of the universal genetic code through the hierarchical nesting of variant codon-amino acid pairings. This evolution was continuously driven by newly distinguishable structural elements of folded proteins being able to distinguish more accurately between amino acids and corresponding tRNA sequence motifs, at each point precisely instantiating a “difference that makes a difference”, which Bateson (1972) defined as the elementary unit of naturally functional information. Although the basic steps taken by nature cannot yet be outlined, we are nonetheless approaching the point where aaRS phylogenetics studies can take us closer to that goal. Furthermore, we can now understand how the self-organised state of coding can be approached “from below”, rather than thinking of it as existing on the verge of a catastrophic fall over a cliff of errors: an incremental improvement in the accuracy of translation will produce replicase molecules that are more faithfully produced from the gene encoding them, probably leading to an incremental improvement in information copying, in turn providing for the selection of narrower genetic quasispecies, an incrementally better encoding of the protein functionalities on which the system relies, including accurate translation. The vicious circle can wind up rapidly from below as a self-amplifying process, rather than winding down the cliff from above, the push-pull tension stably maintaining the system near a tipping point, where, all else being equal, informational replication and translation remain impedance matched – that is, until the system falls into a new vortex of possibilities.

